# Species range expansion constrains the ecological niches of resident butterflies

**DOI:** 10.1101/026765

**Authors:** Hélène Audusseau, Maryline Le Vaillant, Niklas Janz, Sören Nylin, Bengt Karlsson, Reto Schmucki

## Abstract

**Aim:** Changes in community composition resulting from environmental changes modify biotic interactions and affect the distribution and density of local populations. Such changes are currently occurring in nettle-feeding butterflies in Sweden where *Araschnia levana* has recently expanded its range northward and is now likely to interact with the resident species (*Aglais urticae* and *Aglais io*). Butterfly occurrence data collected over years and across regions enabled us to investigate how a recent range expansion of *A. levana* may have affected the environmental niche of resident species.

**Location:** We focused on two regions of Sweden (Skåne and Norrström) where *A. levana* has and has not established, and two time-periods (2001-2006, 2009-2012) during its establishment in Skåne.

**Methods:** We performed two distinct analyses in each region using the PCA-env and the framework described in Broennimann *et al*. (2012). First, we described the main sources of variation in the environment. Second, in each time-period and region, we characterized the realized niches of our focal species across topographic and land use gradients. Third, we quantified overlaps and differences in realized niches between and within species over time.

**Results:** In Skåne, *A. levana* has stabilized its distribution over time while the distribution of the native species has shifted. These shifts depicted a consistent pattern of avoiding overlap between the native species and the environmental space occupied by *A. levana*, and it was stronger for *A. urticae* than for *A. io*. In both regions, we also found evidence of niche partitioning between native species.

**Main conclusions:** Interspecific interactions are likely to affect local species distributions. It appears that the ongoing establishment of *A. levana* has modified local biotic interactions, and induced shifts in resident species’ distributions. Among the mechanisms that can explain such patterns of niche partitioning, parasitoid-driven apparent competition may play an important role in this community.

## INTRODUCTION

Changes in community composition resulting from environmental changes modify biotic interactions that are likely to affect the distribution and the density of local populations. To better predict the widely recorded species’ geographical and environmental shifts (Parmesan, 2006) it is crucial to first define and understand species’ environmental niches and how they are shaped by local biotic interactions (Davis *et al*., 1998; Tylianakis *et al*., 2008; Gilman *et al*., 2010). Investigations of species co-occurrence across the landscape can provide useful insights for better understanding the effects of species interactions on their distributions, and how community composition is maintained or changed locally in a context of global change.

Northern regions offer a highly suitable model system to investigate how changes in species interactions can induce shifts in realized niches. At high latitudes, the impacts of climate change are most pronounced (IPCC, 2014), making it more likely to detect a signature of species’ responses to recent warming. Moreover, as species’ northern ranges vary greatly and their poleward shifts are not synchronized (Pöyry *et al*., 2009), changes in community composition are expected to alter interspecific interactions (González-Megías *et al*., 2008; Devictor *et al*., 2012) and potentially lead to new interactions.

A good way to examine niche shifts and niche partitioning is to measure and test the overlap and the difference in the environmental space occupied (i.e. realized niche) over time and across species (Warren *et al*., 2008; Broennimann *et al*., 2012). Recent developments in statistical methods and computational techniques have enabled better estimations of species-environment relationships and thereby contributed to identify the processes and factors shaping species’ realized niches. In particular, Environmental Niche Models (ENMs) have been used to model and predict species distributions according to changes in climatic and environmental variables, considered to be the main drivers of species distribution at large and small spatial-scales (Berry *et al*., 2002; Thuiller *et al*., 2005). However, ENMs that are based on the relationship between species distribution and abiotic factors have often shown some discrepancy between the potential and the realized niches. This strongly suggests that other processes such as biotic interactions have substantial effect on species distribution and should be accounted for (Leathwick, 1998; Pellissier *et al*., 2012; Tingley *et al*., 2014). Better consideration of biotic interactions in distribution models assessing geographic ranges has greatly refined our predictions at both small and large scales (Heikkinen *et al*., 2007; Araújo & Luoto, 2007). Yet, even though ENMs and ordination techniques have been successfully used to further our understanding of the role of biotic interactions by comparing the realized niches of species (Schweiger *et al*., 2012; Mason *et al*., 2014), they have rarely been used to understand how changes in local biotic interactions may induce shifts in species realized niches (Wisz *et al*., 2013). Furthermore, the reliability and accuracy of statistical models in predicting the importance of biotic interactions are often limited by the availability of extensive occurrence data, which are also often biased towards particular groups of species.

In the last decades, the amount of data collected, organized, and made available through public databases has increased substantially. The use of such databases comes nonetheless with important challenges as they cumulate data collected with no standardized sampling design and by observers with different levels of expertise. Therefore, data contained in such databases have the drawback of being prone to show biases in the region and habitat covered, lacking independence between replicates, and having no explicit measures of sampling effort. However, considering the rapid expansion of programs collating data through volunteer contribution of citizen observers, the real potential of these large datasets is growing and many of them remain largely unexploited. This has led to increased efforts being made to develop robust approaches to estimate and compare the realized niche from occurrence and spatial environmental data, independently of the spatial resolution and sampling biases that are often inherent to species occurrence data. In this context, Broennimann *et al*. (2012) developed and tested an analysis framework to quantify niche overlap and test for niche equivalency and similarity (*cf*. Warren *et al*., 2008), using Principal Component Analysis to define the environmental space (PCA-env). In contrast to ENM methods, the PCA-env has been shown to be more reliable for defining the environmental space when tested on both simulated and real case data (Broennimann *et al*., 2012).

Here, we investigate the realized niche of three nettle-feeding butterflies (*Aglais urticae, Aglais io*, and *Araschnia levana*) and examine how they vary in space and time as an illustration of the potential role of interspecific interactions in shaping species distributions in the context of ongoing climate change. We use occurrence data available through the internet reporting system Artportalen (www.artportalen.se), a public database of species records based on citizens’ contribution in Sweden, to explore niche partitioning of nettle-feeding butterflies at the local scale. In Sweden, *A. urticae* and *A. io* are common native species, easy to identify, and well represented in the database with a large number of records available across the country (over 15 000 records per species for the period 2001-2012). On the other hand, *A. levana*, for which the first anecdotal observation reported in Sweden is from 1982 (Eliasson *et al*., 2005), is known to be expanding its range northward and is now well established in the southern part of Sweden (Betzholtz *et al*., 2013). All three species are specialists with larvae feeding on stinging nettle, *Urtica dioica*, and showing overlapping phenology (see Fig. S1 in Appendix S1 in Supporting Information). While no obvious direct competition has been documented, the three species are known to share common parasitoids (Hinz & Horstmann, 2007; Shaw *et al*., 2009), which increase the potential for apparent competition among the species (van Veen *et al*., 2006; Tylianakis, 2009).

Hence, the ongoing establishment of *A. levana* in southern Sweden makes this system a unique opportunity to investigate the impact of such change on the distribution of native butterfly species over a relatively short time period. If interspecific interactions are important determinants of spatial distribution across the landscape, their signature should be detectable in the occupancy pattern of the available environmental space. To model the realized niches of three sympatric species and quantify the degree of niche overlap between them, we used the PCA-env method applied within Broennimann’s analysis framework (Broennimann *et al*., 2012). We gathered high-resolution data on land use and topography, and an index of nitrogen and phosphorus flows in the soil, which reflects agricultural practices. We perform these analyses over two time-periods during the establishment of *A. levana* and two geographically distinct regions (where *A. levana* has and has not yet established), with the aim of assessing the potential effect of the establishment of *A. levana* in the southern part of the country during the second period, while controlling for climatic variability over time.

Specifically, as a recent colonizer of Sweden, *A. levana* is still in its initial phase of establishment and, therefore, its distribution is expected to not be at equilibrium. Moreover, if changes in local biotic interactions are important in this community, we expect the establishment of *A. levana* to affect the distribution of the native species. We put forward three alternative mechanisms that, in our study system, may explain the observed changes in distribution and density of the local populations. We also discuss the opportunities and limitations of using occurrence records from citizen-science databases and the reliability of the interpretation of the output results from the method used.

## MATERIALS AND METHODS

We studied the environmental space occupied by *A. urticae, A. io*, and *A. levana* in two regions, corresponding to the county of Skåne in Southern Sweden and the Norrström drainage basin in Central Sweden (including four counties: Södermanland, Stockholm, Uppsala, Västmanland, Fig. 1a), and two time-periods (first period 2001-2006 and second period 2009-2012). The regions are separated by approximately 400 kilometers in a straight line. By aggregating the records over specific time-periods, we aimed to have better estimates of species distributions, assuming these patterns to remain relatively stable over short time-periods (i.e. to avoid yearly variation). Data extraction was based on a regular grid covering both regions with a resolution of 1 km.

**Figure 1.**
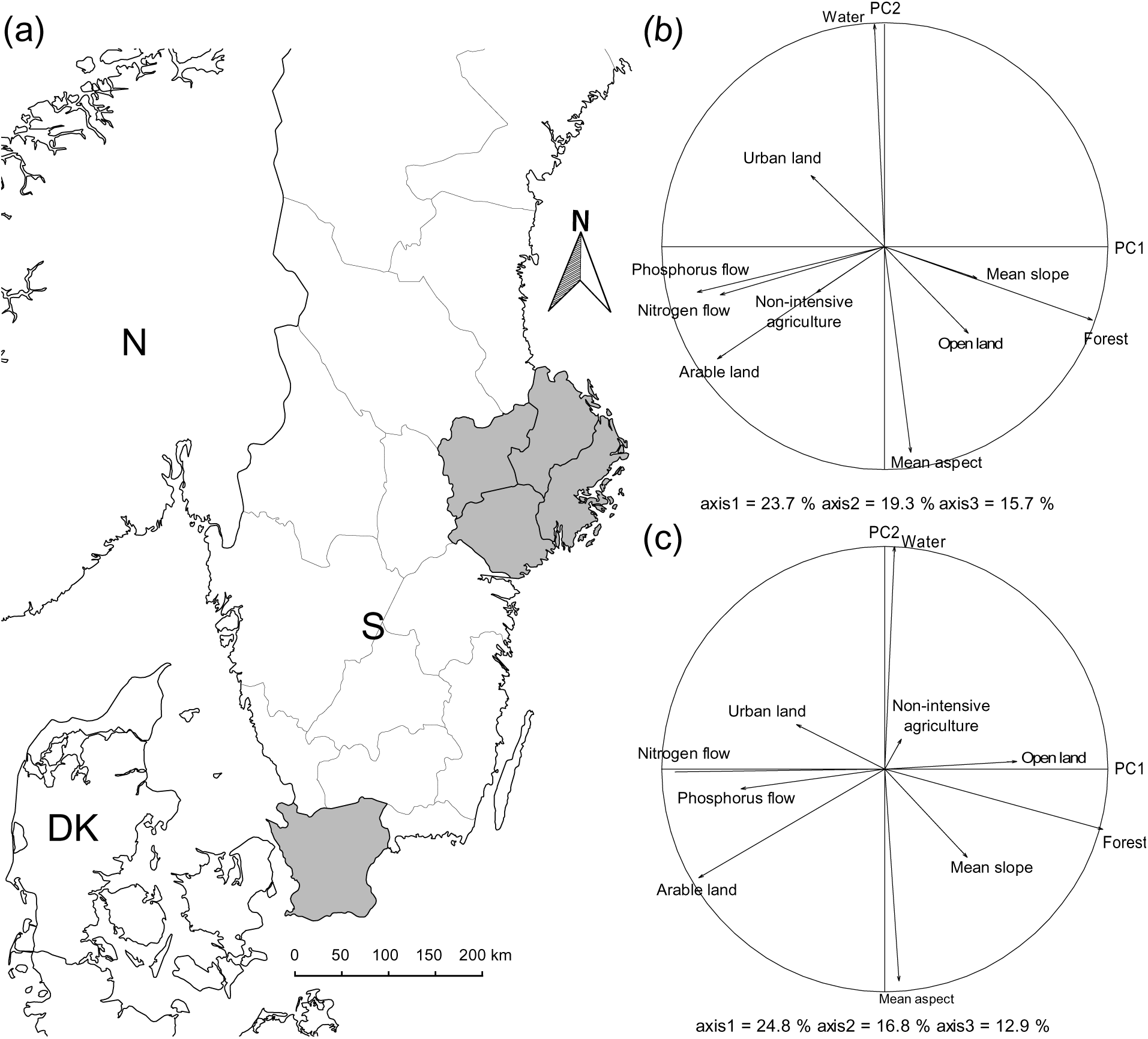
(a) Map of the two study-regions (in grey), corresponding to the county of Skåne in Southern Sweden and the Nörrstrom drainage basin in Central Sweden. (b), (c) Correlation circles showing the contribution of the different variables to the two main axes of the PCA, which describe the environmental space in Nörrstrom and Skåne, respectively. S: Sweden, N: Norway, DK: Denmark.

Species distribution is determined by the interplay of multiple variables operating at different spatial scales (Pearson & Dawson, 2003). While at large scales, species distribution is limited by physiological constraints mainly determined by climatic conditions, at small scales we expect their distribution to be shaped by local variables such as land use and topography (Pearson & Dawson, 2003), as well as by direct and indirect biotic interactions (e.g. predation, competition, resources). Considering the extent of our study areas, i.e. two regions of Sweden, we expected local-scale variables, namely topography and land use, to be the most relevant factors for investigating niche differences between species and potential niche shifts.

### Occurrence Data

We extracted occurrence data for *A. urticae, A. io* and *A. levana* from the Swedish Species Information Centre at SLU (www.artportalen.se, accessed on the 18/09/2015, average precision reported for the data ± se of 168m ± 2m), a public database of species records in Sweden. The Artportalen database gathers opportunistic occurrence data (presence only) collected at 90% by amateurs with no specific required training in species identification (Gärdenfors *et al*., 2014) and who do not follow any specific protocol. For each species, we identify all grid cells for which the species was recorded in each of the two time-periods and the two study regions.

For each period, we only considered species’ presence across the selected grid cells, without accounting for variation in the number of observations per cell as our main interest was to determine species’ occurrence patterns across the total available environment. In this way, we reduced the potential bias associated with uneven sampling effort across sites (i.e. over-sampling of more frequently visited sites). As a result of an overall growing interest for citizen science, both the number of observations and the corresponding number of grid cells visited increased between the first and the second period (Table S1 in Appendix S1). While the number of records is likely to affect the extent of the environmental space sampled (surveyed), we are confident that the sampling (reporting) effort for our study species was sufficiently high to limit this potential bias. For the period 2009-2012, *A. urticae* and *A. io* are the first and fourth most reported butterfly species in Artportalen, respectively, and they were in the top fifteen most reported species according to the Swedish Butterfly Monitoring Scheme (records along transects, Petterson *et al*., 2013). As a recently established species in Sweden, *A. levana* may still be in its initial phase of establishment. Hence, its current distribution might not be at equilibrium yet or only reflect a subset of its potential environmental niche. In our final dataset, occurrence of *A. urticae* was recorded in 2935 grid cells, *A. io* in 2457, and *A. levana* in 599. Note that *A. levana* was only recorded in Skåne.

### Topographic Data

Aspect and slope were calculated based on a digital elevation map at an original 50m resolution obtained from the Swedish University of Agricultural Science (https://maps.slu.se/get/). From the original elevation map, we extracted the mean aspect and mean slope with the r.slope.aspect function available in the GRASS GIS plugin for Quantum GIS 1.8 software (2012) and recalculated both metrics for the 1km resolution grid, using PostgreSQL 9.4 and its spatial extension PostGIS 2.1 (2014).

### Land use Data

Land use data were collected from Naturvårdsverket at an original 25m resolution (https://www.naturvardsverket.se/). The land use classification followed the Corine Land Cover.

We extracted the percentage of the different types of land use (forest, open land, arable land, nonintensive agriculture, water body, and urban land) at 1km resolution grid cells. We also used estimates of soil nutrient flow (in nitrogen and phosphorus) – which we used as a proxy for the amount of nutrients available for plant root absorption – accessible from the SMED (Svenska Miljö Emission Data, http://www.smed.se/) at an original resolution of the sub-catchment (municipality). These estimates were modeled by means of simulation tools as well as measured data and correspond to nitrogen and phosphorus loads to the water from diffuse sources across the whole sub-catchment (Brandt *et al*., 2009). Nutrient loads to the water from the sub-catchment result from a combination of the run-off and leaching on the basis of assumptions on type-specific concentrations for each type of land use. From these estimates, we calculated nutrient flow through the 1km grid cells.

### Realized Niche Shifts and Overlaps

We first extracted two axes that captured the maximum variation in land use and topography available to the species, using the PCA-env ordination method described in Broennimann *et al*. (2012). Second, we characterized, for each time-period and region, the realized niches of our study species in the environmental space defined by the topographic and land use gradients extracted above. Third, we quantified overlaps and differences for each species over time and between species in each period in how they distributed themselves in this environment, using 100% of the environmental space. All analyses were computed with the ecospat package (Broennimann *et al*., 2015) in R 3.1.3 (R Core Team, 2015). To reduce the risk of generating false absences in unsampled areas due to the non-systematic sampling process that characterized our dataset, we excluded from our analyses all grid cells where none of the focal species has been recorded, for each time-period.

Following the framework proposed by Broennimann *et al*. (2012), we computed a weighted Principal Components Analysis (PCA) on the environmental variables, after applying a kernel density function to the number of sites of each specific environmental condition. The Gaussian kernel density function is used to create a probability density function of each of the environmental conditions available and of the occurrence of each species for each cell of the environmental space (Broennimann *et al*., 2012). We performed two distinct analyses in each region to prevent environmental differences inherent to these regions (mostly in agricultural activity) to mask the components of most interest for detecting niche partitioning occurring within region. A kernel density function was also fitted on species occurrence records prior to projecting species occupancy in the environmental space. Thereby, each cell of the environmental space is weighted according to the availability of this specific environmental condition and species occurrence records are weighed in a way that all species involved in the comparison are given similar total weights.

We further tested for niche equivalency and niche similarity (*cf*. Warren *et al*., 2008). For that, we performed paired comparisons of the realized niches of each species over time and between species within each period. The first test evaluates if the environmental conditions that define the niches of two entities are identical. Specifically, niche equivalency is tested by comparing the overlap between the two realized niches with the expected distribution of overlap obtained by randomly reallocating the grid cells occupied by the two entities. The second test assesses the similarity in the relative distribution over the environmental conditions defining the niches of two entities.

Niche similarity is tested by comparing the overlap between the two realized niches to the expected distribution of overlap obtained by reallocating the density of occurrence of one entity across its range of occupancy, while the occurrence of the other remains constant. In other words, this test estimates the likelihood that niche centroids are significantly different from each other. For both tests, expected distributions were based on 500 iterations of the randomization procedure.

## RESULTS

### Description of the available environment and realized niches

The environmental space sampled in each region was described by the first two axes of the principal component analyses, capturing 43.0% and 41.6% of the environmental variation in Nörrstrom and Skåne respectively (Fig 1b & c, and Appendix S2). In both regions, the first PCA-axis was strongly associated to a gradient defined by grid cells having a higher amount of arable land at one end and more forest at the other. Note that in both cases the estimates of soil nitrogen and phosphorus flow explained a large part of the variance captured by the first PCA-axis and were associated with higher amount of arable land. In both regions, the variance along the second axis reflected change in the mean aspect (varying from 0 degree North to 301 degrees at maximum).

Overall, the ecological niches occupied by *A. io* and *A. urticae* in each region between time-periods were highly comparable (Figs 2 & 3).

**Figure 2.**
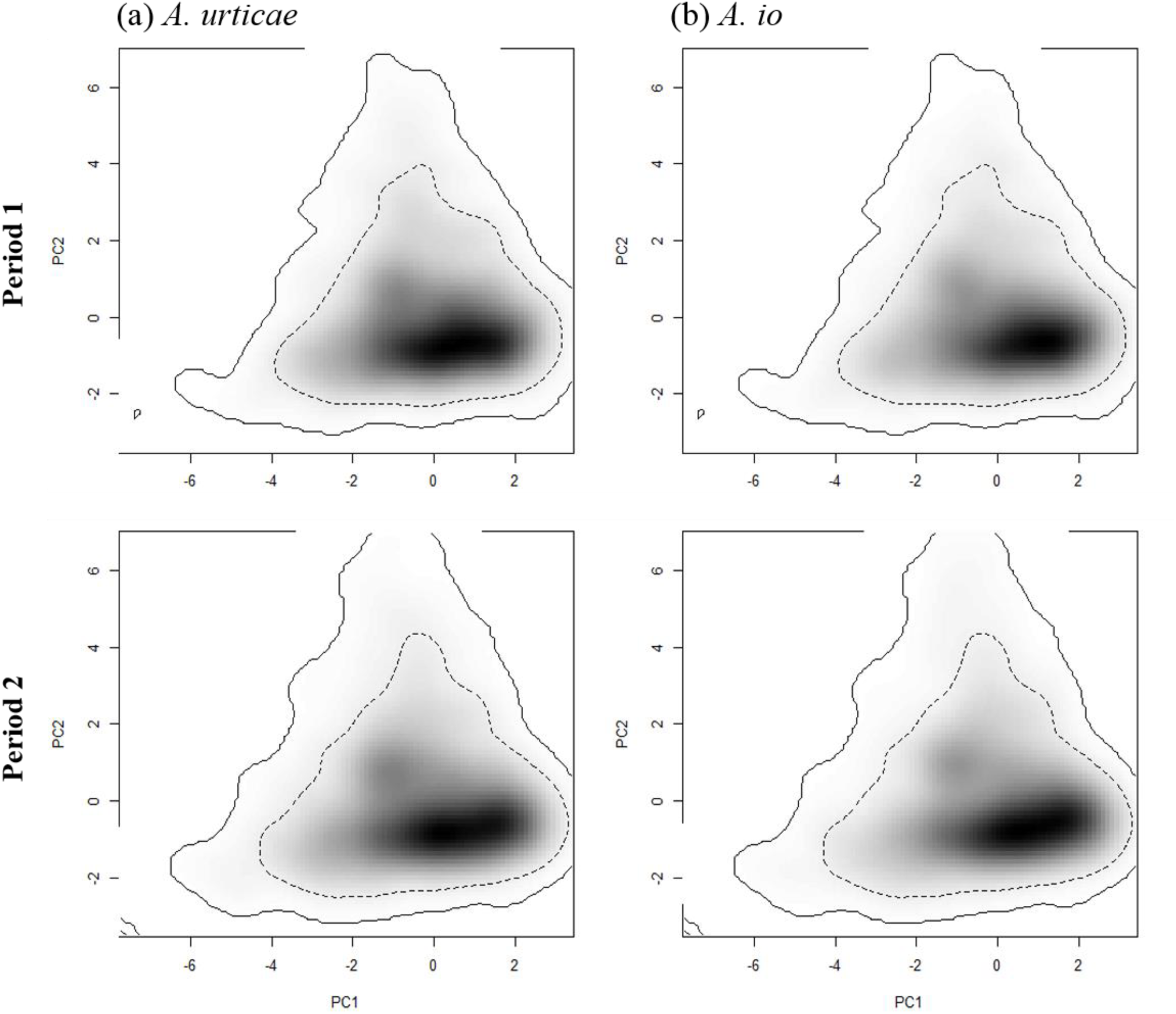
Density of species occurrence across the two-dimensional environmental space describing the Nörrstrom region. In columns are the densities of occurrence of (a) *Aglais urticae* and (b) *Aglais io* for each period (in rows). The black gradient corresponds to the increase in the density of occurrence of the species. The solid line corresponds to the limit of the environmental space available. The dashed line corresponds to the 50% most frequently available environmental conditions.

**Figure 3.**
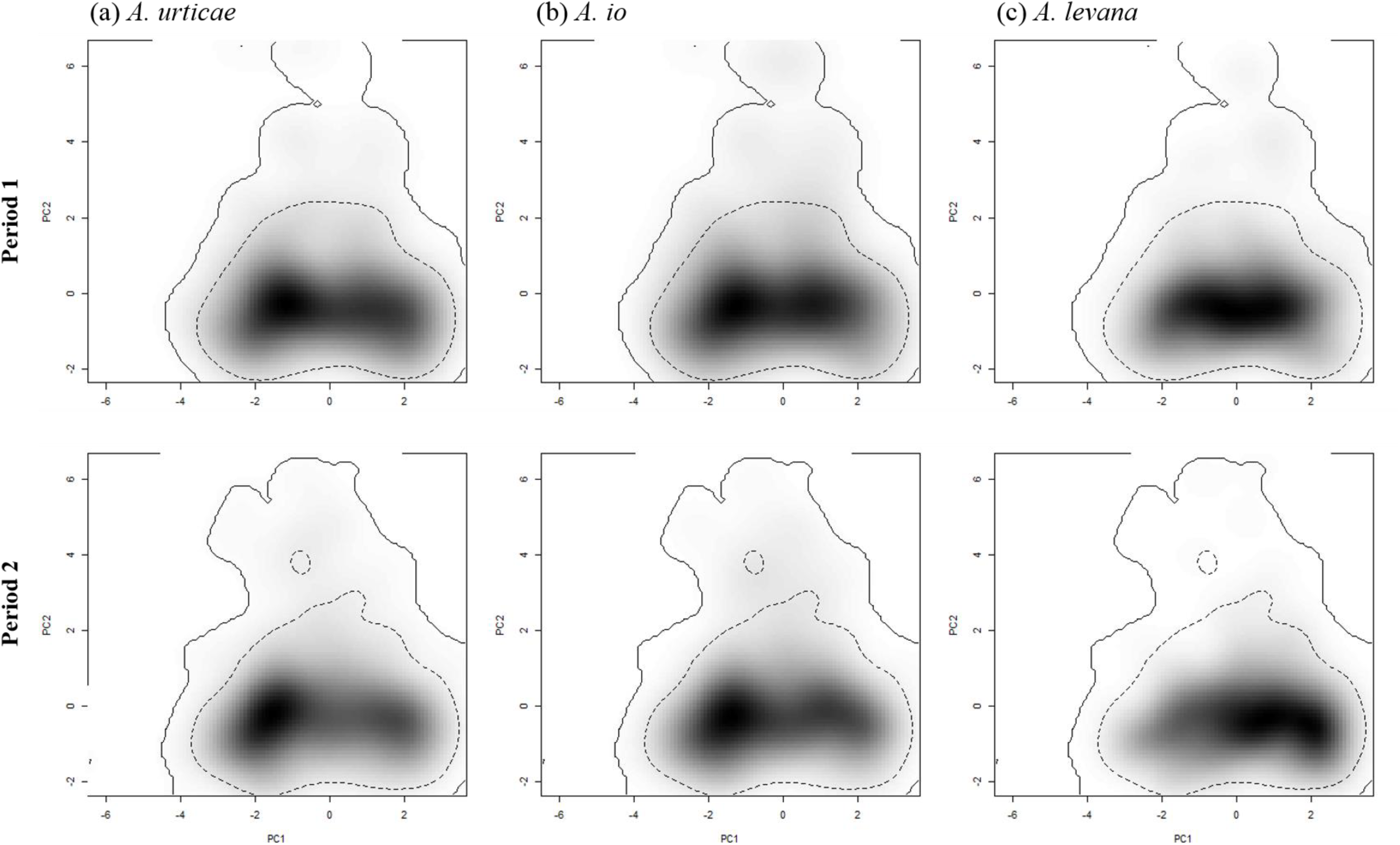
Density of species occurrence across the two-dimensional environmental space describing the Skåne region. In columns are the densities of occurrence of (a) *Aglais urticae*, (b) *Aglais io* and (c) *Araschnia levana* for each period (in rows). The black gradient corresponds to the increase in the density of occurrence of the species. The solid line corresponds to the limit of the environmental space available. The dashed line corresponds to the 50% most frequently available environmental conditions.

### Interspecific interactions

In spite of the overall similarities, we detected in each region and time-period significant differences among the species’ realized niches. In most cases, the null hypothesis of niche equivalency and similarity was rejected (species interactions, *p*<0.05, Table 1), even though the three species displayed important overlap in their environmental space (Table 1, Figs 4).

**Table 1.** Table showing niche overlap (D) within species over time (time shift) and between species in each period (species interactions) in Skåne and Nörrstrom, respectively, with the associated *P*-values obtained from the tests of niche equivalency and similarity (no. iterations=500).

| Region |  | Period | Species | Test of niche equivalency |  | Test of niche similarity |  |
| --- | --- | --- | --- | --- | --- | --- | --- |
|  |  |  |  |  |  | Sp 1 -> Sp 2 | Sp 2 -> Sp 1 |
|  |  |  | Sp 1 - Sp 2 | Overlap D | *p* | *p* | *p* |
| Skåne | Time shift | P1 - P2 | *A. urticae* | 0.82 | 0.004 | 0.002 | 0.002 |
|  |  | P1 - P2 | *A. io* | 0.83 | 0.004 | 0.002 | 0.002 |
|  |  | P1 - P2 | *A. levana* | 0.75 | 0.004 | 0.002 | 0.002 |
|  | Species interactions | P1 - P1 | *A. urticae* - *A. io* | 0.89 | 0.327 | 0.002 | 0.002 |
|  |  | P1 - P1 | *A. urticae* - *A. levana* | 0.80 | 0.004 | 0.002 | 0.002 |
|  |  | P1 - P1 | *A. io* - *A. levana* | 0.80 | 0.004 | 0.002 | 0.002 |
|  |  | P2 - P2 | *A. urticae* - *A. io* | 0.89 | 0.032 | 0.002 | 0.002 |
|  |  | P2 - P2 | *A. urticae* - *A. levana* | 0.68 | 0.004 | 0.002 | 0.016 |
|  |  | P2 - P2 | *A. io* - *A. levana* | 0.68 | 0.004 | 0.002 | 0.004 |
| Norrström | Time shift | P1 - P2 | *A. urticae* | 0.87 | 0.004 | 0.002 | 0.002 |
|  |  | P1 - P2 | *A. io* | 0.85 | 0.004 | 0.002 | 0.002 |
|  | Species interactions | P1 - P1 | *A. urticae* - *A. io* | 0.91 | 0.032 | 0.002 | 0.002 |
|  |  | P2 - P2 | *A. urticae* - *A. io* | 0.89 | 0.004 | 0.002 | 0.002 |

**Figure 4.**
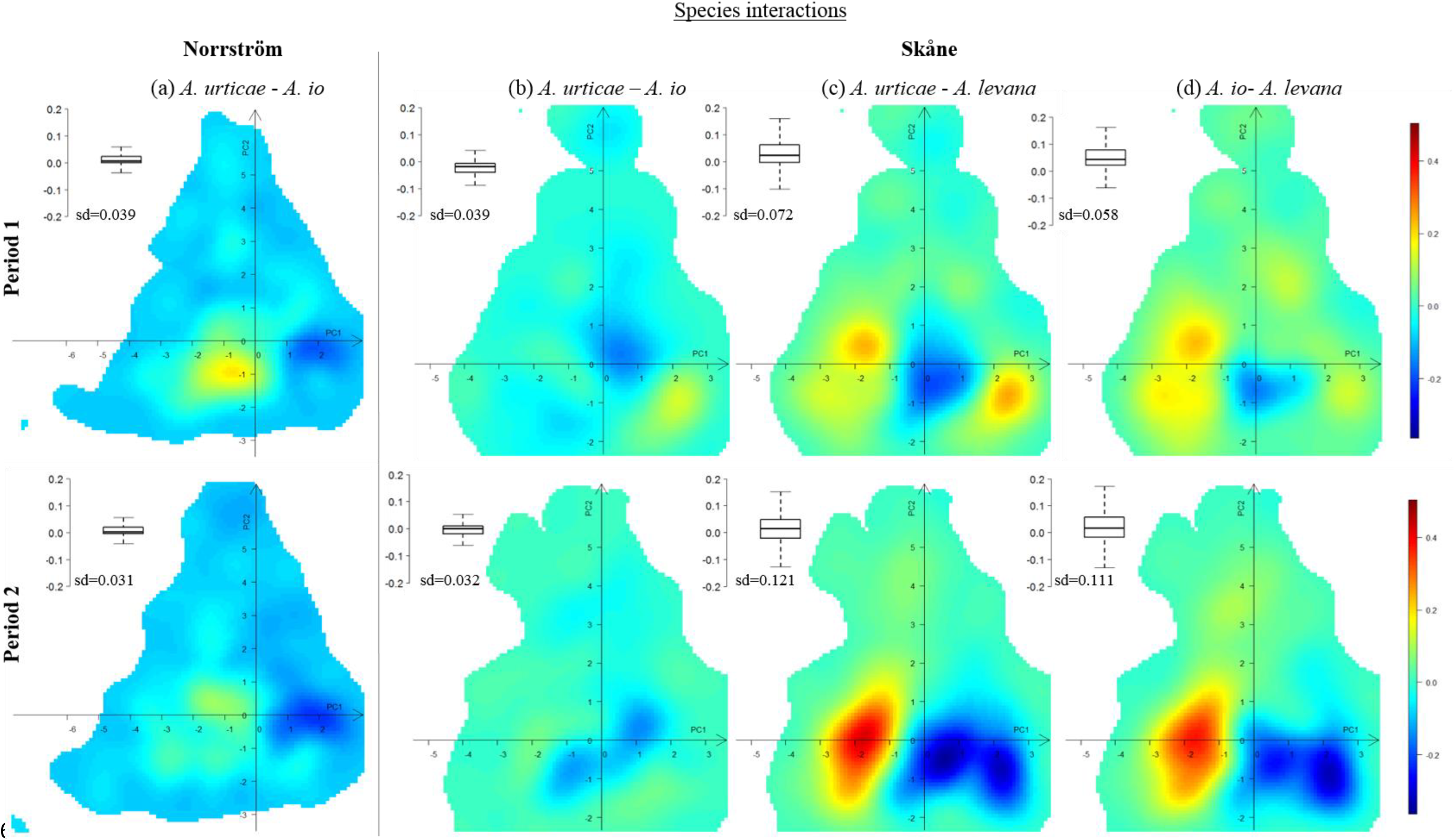
Plots of the differences in density of occurrence, for each region and time-period, between species. In columns are represented the differences in the density of occurrence in Nörrstrom of (a) *Aglais urticae* and *Aglais io* and in Skåne of (b) *A. urticae* and *A. io*, (c) *A. urticae* and *A. levana* and (d) *A. io* and *A. levana*, for each period (in rows). Red cells indicate the prevalence of the first species over the second species. Dark blue cells indicate the prevalence of the second species over the first species. Grid cells with a color value of zero correspond to environmental conditions for which the densities of occurrence of both species were similar. The boxplot and the standard deviation (sd) associated with each graph show the variability of the differences of densities observed across the available environment. The larger the box and the higher the sd are, the more contrasting the species are in their distribution.

In the two-dimensional environmental space in Skåne, *A. levana* overlapped with the two native species in both time-periods but the species’ realized niches were neither equivalent nor similar (*p*<0.05, Table 1). Moreover, we observed a large decrease in the overlap of *A. levana* with the native species over time. In the first period it was 0.80 with *A. urticae* and *A. io*, while in the second period it was 0.68 with both native species. In the first period we did not observe a clear differentiation between *A. levana* and the resident species’ niches (Fig. 4c & d top), but in the second period this partitioning appears clearly (Fig. 4c & d bottom). In the second period, *A. levana* preferentially occupied habitats with larger amount of forest per 1 km grid cell and the two native species were in comparison more present in mixed and agricultural habitats (Fig. 4c & d bottom).

The two native species (*A. urticae* and *A. io*) strongly overlapped in both periods and regions (overlap between 0.89 and 0.91, Table 1). This is not surprising considering that *A. io* and *A. urticae* are often observed in sympatry with ecological niches that are known to be difficult to differentiate. While in the first period, *A. urticae* and *A. io* displayed equivalent niches in Skåne (paired-species comparisons of niche equivalency, *p*=0.327, Table 1), the null hypothesis of niche equivalency was rejected in the first period in Nörrstrom and in both regions in the second period *p*<0.05, Table 1). Thus, although the realized niches of *A. urticae* and *A. io* significantly overlapped, they displayed non-random differences in their distributions. For all the paired-species comparisons between *A. urticae* and *A. io*, we rejected the null hypothesis of niche similarity (tests of niche similarity, *p*<0.05, Table 1); which suggests that these species occupied the available environment differently in both time-periods and regions (shifts in their centroid). In Skåne in the first period, *A. urticae* occupied proportionally more areas with larger amount of forest per 1 km grid cells than *A. io*, whereas *A. io* was more abundant in mixed habitats (Fig. 4b top). In the second period, the prevalence of *A. urticae* slightly shifted in comparison to *A. io* from forested areas toward habitats with increasing amount of agricultural and urban lands (Fig. 4b bottom). In Nörrstrom and for both periods, *A. urticae* and *A. io* prevailed in environments with higher amount of forest (Fig. 2).

### Niche shifts over time

The distribution of the three species shifted significantly over time in both regions, their distribution in the environmental space being neither equivalent nor similar between the two time-periods (tests, *p*<0.05, Table 1). The largest shift was observed for *A. levana* in Skåne, for which the overlap in the realized niche between periods was of 0.75. For the two native species these shifts were comparatively smaller, the overlap in the realized niche of each species between periods varied between 0.82 and 0.87 (Fig. 5), but noticeable considering the short time scale of the study.

**Figure 5.**
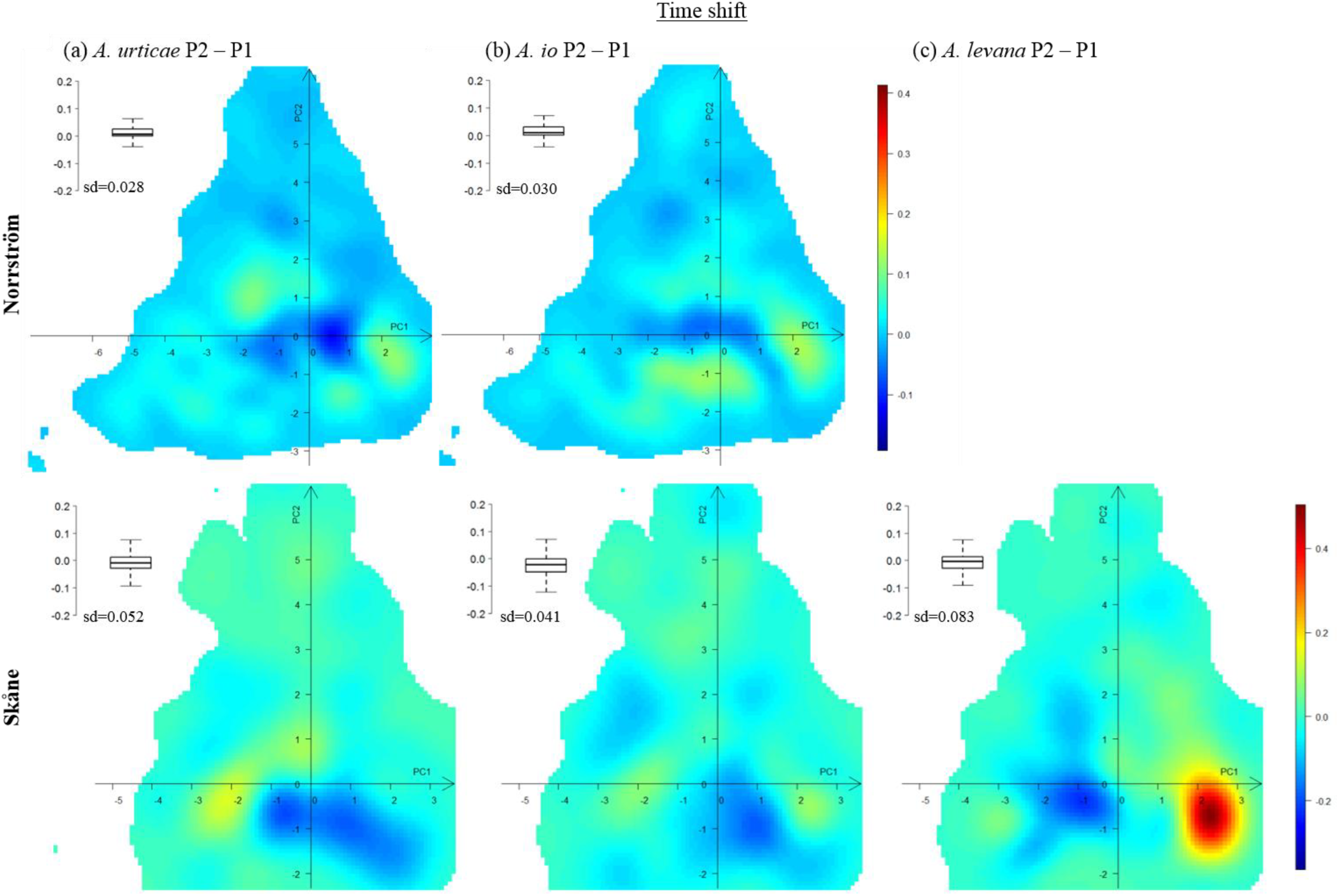
Plots of the differences in density of occurrence of each species over time, and for each region. In columns are represented the differences in the density of occurrence of (a) *Aglais urticae*, (b) *Aglais io* and (c) *A. levana* over time and for the regions Nörrstrom and Skåne (in rows). Red cells indicate the prevalence of the species in the second period. Dark blue cells indicate the prevalence of the species in the first period. Grid cells with a color value of zero correspond to environmental conditions for which the density of occurrence of the species did not vary over time. The boxplot and the standard deviation (sd) associated with each graph show the variability of the differences in densities observed across the available environment. The larger the box and the higher the sd are, the more contrasting the species distributions are.

All shifts detected in the environmental space also corresponded to geographical shifts, as only one layer of land use was available to cover both time-periods. For the recently established *A. levana*, the observed shift in realized niche corresponded to a niche contraction as *A. levana* in the first period seemed to have already colonized most of the available environmental conditions in which the species occurred in the second period (Fig. 3c). In the second period, *A. levana* preferentially occupied areas with larger amount of forest (Fig. 5c bottom). For the two native species, the largest changes between the two periods were observed in Skåne (Fig. 5a & b). Moreover, the extent of the environmental shift observed in Skåne was species-specific, being larger for *A. urticae* (niche overlap=0.82, sd=0.052; Fig. 5a bottom) than for *A. io* (niche overlap=0.83, sd=0.041; Fig. 5b bottom). The shift was also more clearly directed for *A. urticae* than for *A. io* (Fig. 5a & b bottom). By shifting its distribution away from areas with larger amount of forest, *A. urticae* reduced its overlap with *A. levana*, which over time has densified its distribution in these environments across Skåne (niche overlap=0.68, sd=0.121; Fig. 4c bottom). We observed the same pattern between *A. io* and *A. levana*, but to a lower extent (niche overlap=0.68, sd=0.111; Fig. 4c bottom).

## DISCUSSION

The distributions of our focal species were described and compared along the two first axes of the environmental PCA. In both regions, the first axis was shaped along a gradient from arable land to more forested land and the second axis correlated with topographic parameters (Fig.1a & b). Based on our knowledge of the ecological niche of our focal species, we are confident in the description of the species’ environmental niche revealed by our models. The method developed by Broennimann *et al*. (2012) gave results consistent with the overall characteristics of the realized niches of the three butterfly species, which are known to occupy open habitats and woodlands, wood hedges, and hedgerows (Asher *et al*., 2001; Haahtela *et al*., 2011).

Yet, the method also identified differences in realized niches among species in each region and within species over time. We used the measure of overlap and the results from the niche equivalency and similarity tests between realized niches to infer the role of interspecific interactions in shaping species distributions. The large overlap in the realized niches of the two native species in both regions and time periods suggests that no clear negative interaction is involved in the partitioning of the niche between *A. urticae* and *A.io*. Nevertheless, the fact that the realized niches of species were neither equivalent, nor similar, suggests some level of nonrandom niche partitioning in the environmental space. Thus, while our results emphasized that the two native species occupy the available environment differently, we could not identify a specific environmental factor of niche differentiation. For example, we have observed in the field that *A. urticae* tends to occupy more sun-facing slopes than *A. io*, but although we detected some effect of aspect on species’ distribution, the relatively coarse resolution of our data (1km grid cell) prevents us to firmly support our field observations. Moreover, apart from the spatial component, niche partitioning can also arise from, and in combination with, resource and/or temporal partitioning (MacArthur & Levins, 1967). Regarding resource partitioning, both *A. urticae* and *A. io* larvae feed on the leaves of stinging nettles, whose availability is not limited over the season. Stinging nettle has even been shown to expand due to human land use and nitrogen pollution (Taylor, 2009). Together with the lack of strong competition over resources, potential asynchrony (temporal partitioning) in species phenology could also reduce the strength of a spatial signal in the differentiation in species’ realized niches.

The recently established *A. levana* overlaps with the realized niches of the native species and the overlap decreased over time. In part, this decrease is likely a consequence of the described shifts in realized niche of the three species over time. The largest shift was found for *A. levana* (in Skåne). From the environmental conditions colonized by *A. levana* in the first period, we observed a significant contraction in its realized niche in the second period toward habitats with a higher amount of forest. This pattern suggests that the distribution of *A. levana* was not at equilibrium in the first period and that it has progressively stabilized within habitats with a positive population dynamic. We also found significant shifts in realized niches of the two native species over time, in both regions. However, these shifts were most pronounced in Skåne where *A. urticae*, and to a lower extent *A. io*, shifted their distributions away from forested areas towards habitats with higher amount of agricultural and urban lands.

There are several processes that could explain these changes over time. The more pronounced shift in the distribution of the native species in Skåne than in Nörrstrom could partly reflect higher regional climatic fluctuations over time. However, as the amplitudes of climatic variations over time were comparable across the regions (Appendix S3), this factor does not appear to be a major driver in our study system. The observed shifts in niche centroid over time could also be related to the increase in the number of reported observations through time. Yet, this explanation also appears rather unlikely, as the increases in the number of observation records were consistent across species and regions, while all species did not show a consistent shift in their centroids. Thus, even if such shifts may have been strengthened by the higher number of observation records over time, they most likely represent genuine responses to other driving forces affecting the ecological niche of native species.

Hence, in this community, changes in interspecific interactions appear to be a sensible explanation for the described changes over time in species occupancy of the environmental space. One contributing factor is probably the stabilization of the distribution of *A. levana*, as this resulted in decreased overlap with the native species (Fig. 4c & d). This response may have been mediated through the action of parasitoids (Dunn *et al*., 2012). Indeed, studies have shown that the community composition in herbivorous insects can largely be shaped by parasites (van Veen *et al*., 2006; Tylianakis, 2009). All three species are heavily parasitized (personal observation) and share many parasitoids (Hinz & Horstmann, 2007; Shaw *et al*., 2009). Hence, the population dynamics of these parasitoids might have been positively affected by the increase in potential hosts (*A. levana*) between the two time-periods, as observed in tropical forest communities by Morris *et al*. (2004). Moreover, the differences in phenology between *A. urticae, A. io*, and *A. levana* (Fig. S1 in Appendix S1) may form a continuous breeding ground to stimulate parasitoid population dynamics, and they may result in a prolonged temporal niche for parasitoids (Blitzer & Welter, 2011). In a scenario of parasitoid-driven apparent competition, the phenologically late species is expected to be the more vulnerable as its life cycle will coincide with an increase in parasitoid population size (Blitzer & Welter, 2011). In this study, the native species showed differences in the magnitude of their shifts. The shift in the realized niche of *A. urticae* between periods was stronger than for *A. io*, in terms of niche overlap. Niche partitioning was also stronger between *A. levana* and *A. urticae* than with *A. io*. In the context of parasitoid buildup, this may seem counterintuitive, as *A. urticae* is the earliest of the two native species. But because *A. urticae* is bivoltine in Skåne, and produces a second brood later in the season (Fig. S1 in Appendix S1), parasitoid populations may continue to build up over the season and result in the highest parasitoid load for the second brood of *A. urticae*. Although we have no direct data to support the hypothesis of parasitoid-driven niche differentiation, the circumstantial evidence is suggestive, and is further reinforced by the observation that *A. levana* appears to share more parasitoids with *A. urticae* than with *A. io*, based on the limited information available (Shaw *et al*., 2009).

Differences in life history might be another possible explanation for the different shifts in distribution among the three species. Based on experimental studies, Merckx *et al*. (2015) suggested that because of its bivoltine life cycle, the plastic response of *A. urticae* to anthropogenic environmental changes might be faster in comparison to the univoltine *A. io*. If this is true, *A. urticae* would be expected to respond faster to the arrival of *A. levana* than *A. io*. This is consistent with the described larger shift in distribution of *A. urticae* than *A. io* over time in Skåne. Likewise, the bivoltine life cycle of *A. levana* may also participate to explain the large shift described in its distribution over time.

It is also possible that differences in the way these species are able to adjust to the quality of their host plants and the microclimate they offer may cause individuals of each species to choose different nettle patches when coexisting in a region. The shift in niche centroid of *A. urticae* reported here (toward agricultural land) is consistent with other studies conducted on *A. urticae* and *A. io*, showing that *A. urticae* can to a larger extent be favored by the higher nutrient quality of plants growing in agricultural land than *A. io* (Serruys & Van Dyck, 2014; Audusseau *et al*., 2015; Merckx *et al*., 2015). Changes in microclimate have been shown to differentially affect the decline of butterfly species’ following climate change (Wallisdevries & van Swaay, 2006) and to modify butterflies’ habitat associations (Davies *et al*., 2006). We call for field experiments to understand, at a finer spatial-scale, the differences among species in their optimal microclimatic niche.

The method used was successful in handling the potential bias related to the non-standardized sampling design associated to occurrence data extracted from the public database. Indeed, while most observations were recorded in urban areas, the highest densities of occurrence of the two native species based on the grid cell analysis were found in forest habitats (Figs 2 & 3). In addition, the lower number of species records in the first period did not seem to have affected our capacity to extract the main components of the realized niches as their shapes and centroids were consistent through time. Hence, we consider that using occurrence records from citizen-science databases offer great opportunities for investigating the realized niches of species and differences between them, as long as the biological relevance of the modelling results is carefully considered. The results of the method we used to characterize the realized niches of the focal butterfly species was consistent with our knowledge of the species, and we recommend using occurrence data associated with species and regions of high sampling effort in order to get more robust predictions. Such regions often correspond to more densely populated areas as the reporting effort in citizen-based monitoring programs is strongly related to the number of participants.

In conclusion, we found indications of niche partitioning among the three species and consider that the observed shifts in species environmental niche were partly driven by the recent changes in interspecific interactions. The establishment during range expansion of *A. levana* may have modified biotic interactions, resulting in associated shifts in species distributions. Based on our knowledge of the system, we suspect that the observed niche differentiation might be partly driven by apparent competition mediated by shared parasitoids. Other factors such as interspecific differences in life history could also have contributed to the observed pattern and lead to species-specific change in habitat preference. The difference in the extent of the shift observed between *A. urticae* and *A. io* during the establishment of *A. levana* can be related to their differences in phenology, voltinism, and the higher number of shared parasitoids between *A. urticae* and *A. levana*. Further investigations of the niche partitioning of these three nettle-feeding butterflies where they all are native species would allow us to make predictions about the equilibrium this community may reach. More importantly, this framework is a promising tool for investigating the potential role of biotic interactions on species distributions, and for better predicting the outcome of their modifications on community composition and species-environment relationships.

## ACKNOWLEDGMENTS

We thank the Strategic Research Program Ekoklim at Stockholm University for funding of this project. HA is acknowledging support from Helge Ax:son Johnsons stiftelse, RS from the FRB and EDF SA (FRB-CESAB project LOLA-BMS), and SN from the Swedish Research Council (2011-5636). We also wish to thank Vlad Dincã and Claudie Daniel for the photos presented in the Supporting Information, and Lars B. Pettersson for an insightful comment on the manuscript. We are also thankful for the comments from C. Stefanescu and two anonymous referees.

## SUPPORTING INFORMATION

**Appendix S1** Data description, phenology of the study species and consideration of biases.

**Appendix S2** Table showing scores for each variable on the first three axes of the PCA-env and the contribution of each axis expressed in percentage of the total variance (inertia).

**Appendix S3** Average mean monthly temperature (Jan: 1, Feb: 2, etc.) in Skåne in comparison to Nörrstrom and in the first and second periods (mean±se).

**BIOSKETCH** Hélène Audusseau has been granted her PhD at Stockholm University. Her research is concerned by the ecological and evolutionary responses of nettle-feeding butterflies to changes in climate and land use.

**Editor:** Daniel Chapman.

## AUTHORS’ CONTRIBUTIONS

AH, ML, RS, NJ, SN participated in the design of the study. AH, ML, RS performed the analyses. AH, ML, BK collected the data. AH wrote the first draft of the manuscript, which was substantially improved through the contribution of RS, ML, NJ. All authors contributed to the final version of the manuscript.

